# Population size interacts with reproductive longevity to shape the germline mutation rate

**DOI:** 10.1101/2023.12.06.570457

**Authors:** Luke Zhu, Annabel Beichman, Kelley Harris

## Abstract

Mutation rates vary across the tree of life by many orders of magnitude, with lower mutation rates in species that reproduce quickly and maintain large effective population sizes. A compelling explanation for this trend is that large effective population sizes facilitate selection against weakly deleterious “mutator alleles” such as variants that interfere with the molecular efficacy of DNA repair. However, in multicellular organisms, the relationship of the mutation rate to DNA repair efficacy is complicated by variation in reproductive age. Long generation times leave more time for mutations to accrue each generation, and late reproduction likely amplifies the fitness consequences of any DNA repair defect that creates extra mutations in the sperm or eggs. Here, we present theoretical and empirical evidence that a long generation time amplifies the strength of selection for low mutation rates in the spermatocytes and oocytes. This leads to the counterintuitive prediction that the species with the highest germline mutation rates per generation are also the species with most effective mechanisms for DNA proofreading and repair in their germ cells. In contrast, species with different generation times accumulate similar mutation loads during embryonic development. Our results parallel recent findings that the longest-lived species have the lowest mutation rates in adult somatic tissues, potentially due to selection to keep the lifetime mutation load below a harmful threshold.

**Significance Statement:** All cells accumulate mutations due to DNA damage and replication errors. When mutations occur in germ tissues including sperm, eggs, and the early embryo, they create changes in the gene pool that can be passed down to future generations. Here, we examine how rates of germline mutations vary within and between mammalian species, and we find that species which reproduce at older ages tend to accumulate fewer mutations per year in their sperm and eggs. This finding suggests that the evolution of humans’ long reproductive lifespan created evolutionary pressure to improve the fidelity of DNA maintenance in germ tissues, paralleling the pressure to avoid accumulating too many mutations in the body over a long lifespan.

## Introduction

Germline mutation rates vary by orders of magnitude across the tree of life and ultimately limit the adaptability and the complexity of each species (1–5). Low mutation rates may limit the rate of adaptation to new challenges (6–8), while high mutation rates may limit the ability of a well-adapted population to maintain its fitness and dominance (9, 10). Maintenance of a low mutation rate also incurs an energetic cost, requiring investment of resources and genomic real estate in DNA repair machinery and other mutation-avoiding systems (11–14). As organisms get more complex, the possible consequences of a high mutation rate get more complex as well, leading to confusion and debate about which evolutionary forces ultimately shape this important parameter (15–17).

One widely cited model, the drift barrier hypothesis, posits that mutation rate variation is largely driven by differences in effective population size that modulate the efficacy of selection against weakly deleterious alleles (5, 18–20). A “mutator allele” that raises the germline mutation rate is likely to be deleterious given that harmful mutations outnumber beneficial mutations, but since most mutations are neutral or only weakly harmful, a modest increase in the mutation rate is only expected to decrease fitness by a small amount (21, 22). A corollary of the drift barrier hypothesis is that genetic drift likely limits the ability of DNA repair enzymes to function near their biophysical optima, since optimal functioning would require natural selection to weed out mutator alleles that cause very few additional germline mutations each generation and thus have nearly-neutral fitness effects (23). As a result, different nearly-neutral mutator alleles are likely to accumulate over time in each population and species, causing the molecular efficacy of each DNA repair enzyme to diverge across the tree of life (24, 25). Although there exists little direct data on the molecular efficacy of DNA repair and how it varies among species, the predictions of the drift-barrier hypothesis enjoy broad indirect support from mutation rate data, which are easier (though still expensive) to measure. Across the tree of life, population size is inversely correlated with the mutation rate per site per generation (26), and a similar correlation was recently measured using vertebrate mutation rate data alone (27).

In single-celled organisms, there is a fairly direct connection between DNA repair efficacy and mutation rate per generation (which is the same as the mutation rate per cell division). Single-celled organisms also exhibit substantial diversity in the architecture of DNA repair, ranging from the minimalist repair systems of some obligate symbionts (which have very high mutation rates (28)) to unique genomic proofreading mechanisms in ciliates such as *Paramecium*, which have some of the lowest mutation rates known to science (29–31). In contrast, multicellular eukaryotes have more standardized cellular housekeeping processes but varied, multi-stage life histories, with each generation involving multiple cell divisions as well as potentially mutagenic cell states associated with sex and embryonic development (32–34). This complexity muddies the relationship between the mutation rate and the molecular efficacy of DNA repair and complicates the interpretation of the correlation between mutation rate and effective population size. When Bergeron et al. noted that effective population size was correlated with mutation rate among vertebrates, they noted that a similar amount of vertebrate mutation rate variation could be explained by generation time: the typical interval between reproduction events (27). A strong negative correlation between generation time and the mutation rate per generation was previously inferred from phylogenetic substitution data, and the etiology of this pattern has been long debated (16, 17, 35). Measurements of mutation rate variation within human families have made it clear that generation time can influence the mutation rate independently of molecular DNA repair efficacy: as parents age, their children are born with more and more mutations (36, 37).

The effect of parental age on the human mutation rate has been well characterized thanks to the availability of thousands of mutation rate measurements from trios where the ages of the parents at the birth of the child are known (38). Similar (though smaller) trio datasets have also been generated for several non-human mammalian species, and all show the same qualitative pattern of increasing mutation rate as a function of parental age (39–45). These data show evidence of significant mutation rate differences among species, and they also differ in estimates of the rate at which mutation rates increase with the ages of the father and mother. However, the same sample sizes of most non-human mutation rate studies come with high degrees of statistical uncertainty, and some recent studies of mutation rates in primates and carnivores have argued that parental age effects in these species are not statistically distinguishable from each other (40, 44). Instead, they found that mutation rate measurements from several primate species, as well as the domestic cat, were consistent with a *reproductive longevity model* where the molecular efficacy of DNA repair is assumed to be invariant among species and mutation rate differences are instead driven by differences in the timing of puberty and reproduction.

Here, we study the etiology of vertebrate mutation rate variation by decomposing it into its three main components: the rate of mutations that accumulate during embryonic development, the rate of mutations occurring in the gametes per year of adult reproductive life, and the length of the time elapsed between puberty and reproduction. Embryonic and gamete mutation rates are molecular parameters that reflect rates of DNA damage and repair in two very different germ tissues, while the time elapsed between puberty and reproduction is a demographic parameter that varies due to a combination of biology and environmental contingency. Extending the theoretical framework of the drift-barrier hypothesis, we separately model the fitness effects of variation in the embryonic and gamete mutation rates and infer that the fitness effects of alleles that increase the gamete mutation rate are likely to scale with generation time. This scaling reverses the direction of one drift-barrier hypothesis prediction, implying that selection against gamete mutator alleles will be most effective in species with long generation times, not in species with large effective population sizes that tend to have short generation times. We test our predictions by estimating gamete and embryonic mutation rates from published regressions of mutation rate against generation time from eight mammalian species: consistent with our model, we find that generation time appears to be positively correlated with the embryonic mutation rate but negatively correlated with the gamete mutation rate. We go on to show that mutation rate variation among species is broadly consistent with a “relaxed clock” reproductive longevity model where embryonic mutation rates vary according to the classic drift-barrier hypothesis predictions but gamete mutation rates are shaped by a modified drift-barrier model where selection against mutators is intensified by late reproduction.

## Results

### As generation time increases, mutation rates in the embryo and the gametes trend in opposite directions

Several recent papers have modeled the etiology of germline mutations by first separating mutations occurring in early development from mutations that occur post-puberty in the parents’ germ cells (40, 46, 47). In this context, Thomas et al. proposed that there is little variation among mammals in the total mutation load occurring before puberty and in the mutation rate per year occurring in the gametes after puberty, but that most germline mutation rate variation is caused by variation in two demographic parameters: the age of puberty and the time elapsed between puberty and reproduction. To paraphrase the mathematical description of their model, we will let *P* denote the age of puberty, *A*_*M*_ and *A*_*P*_ denote maternal and paternal ages at conception of an offspring, *μ*_*E*_ denote the rate per generation of mutations that accumulate in the embryo before puberty, and *μ*_*S*_ and *μ*_*O*_denote the mutation rates per year in mature spermatocytes and oocytes. In terms of these variables, the germline mutation rate *u*_*g*_ as a function of parental age is:

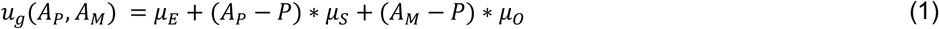

When Thomas, et al. and Wang, et al. published mutation rate data for owl monkeys (40), rhesus macaques (42), and domestic cats (44), they inferred species-specific values of the mutation rate parameters *μ*_*E*_ and *μ*_*O*_ + *μ*_*S*_ (relatively few germline mutation rate studies currently have the power to infer *μ*_*O*_ and *μ*_*S*_ separately). However, they also argued that these species-specific values did not fit the mutation rate data significantly better than a unified model that uses mutation rate parameters *μ*_*E*_, *μ*_*O*_, and *μ*_*S*_ that were previously inferred from human mutation data. **Figure 1A,B** illustrates how this reproductive longevity model can explain variation in mutation rates between species, while **Figure 1C** illustrates a contrasting model where mutation rate variation is driven by variation in the rate parameters *μ*_*E*_ and *μ*_*O*_ + *μ*_*S*_ .

**Figure 1:**
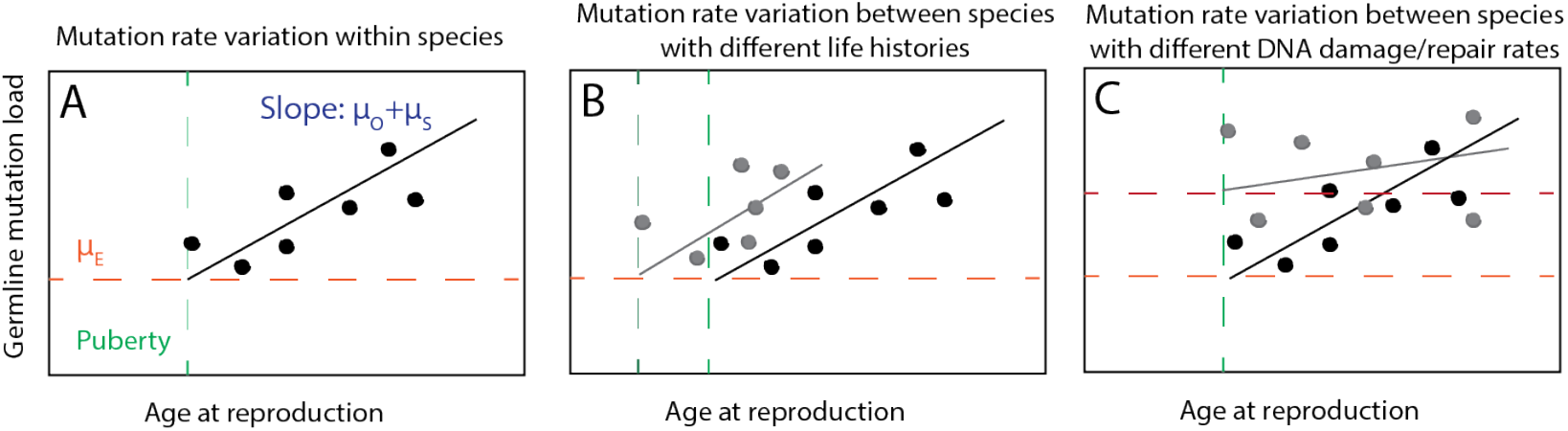
Models of germline mutation rate variation. **A**. Within species, mutation rates vary as a function of age at reproduction. Each individual is expected to accumulate an embryonic mutation load *μ*_*E*_ plus inherit mutations that accumulated in their parents’ sperm and eggs at rate *μ*_*O*_ + *μ*_*S*_ each year between puberty and conception. **B**. Two species with different lifespans and/or different ages of puberty onset may have different distributions of mutation rates despite similar mutation parameters *μ*_*E*_ and *μ*_*O*_ + *μ*_*S*_, as posited in (40, 44). **C**. Two species with similar lifespans and similar ages of puberty onset might still have different mutation rates due to genetic differences that affect rates of DNA damage, repair, or proofreading. This type of mutation rate variation is driven by variation in the parameters *μ*_*E*_ and/or *μ*_*O*_ + *μ*_*S*_ .

Although the constant-rate reproductive longevity model appears to explain much of the mutation rate variation among humans, owl monkeys, macaques, and domestic cats, Lindsay, et al. 2019 previously noted that the spermatocyte mutation rate per year was 5-fold higher in mice compared to humans (48). To formally test whether the Lindsay, et al. mouse data reject a constant-rate reproductive longevity model, we inferred *μ*_*E*_ and *μ*_*O*_ + *μ*_*S*_ from the Lindsay, et al. mouse DNM dataset. We found that these rate parameters both significantly diverged from their human counterparts, with disjoint 95% confidence intervals. In mice, the embryonic mutation rate *μ*_*E*_ = 3.75 × 10^−9^ (95% CI 2.89 × 10^−9^; 4.6 × 10^−9^), while in humans the rate is nearly 2-fold higher: *μ*_*E*_ = 6.35 × 10^−9^ (95% CI 5.47 × 10^−9^; 1.21 × 10^−8^). Conversely, the mouse gamete mutation rate *μ*_*O*_ + *μ*_*S*_ = 1.64 × 10^−9^ (95% CI 4.10 × 10^−10^; 2.85 × 10^−9^), while in humans, the rate is 5-fold lower, as previously noted: *μ*_*O*_ + *μ*_*S*_ = 3.5 × 10^−10^ (95% CI 3.3 × 10^−10^; 3.7 × 10^−10^).

To test whether the difference between mouse and human mutation rate parameters is representative of a broader dependence of these rate parameters on generation time, we searched the literature for other regressions of mutation rate against parental age that would permit estimation of *μ*_*E*_ and *μ*_*O*_ + *μ*_*S*_ for additional species. We found appropriate data for five additional primates plus two carnivores, transformed these species-specific regression parameters into standardized mutation rate units, and compiled these parameters in **Table 1**.

**Table 1:** Regression-based estimates of embryo and gamete mutation rates. The generation times and ages at first reproduction in the table are drawn from the publications reporting each set of mutation rate data. See Supplementary Methods for a description of how these standardized rates were calculated from each study’s reported data.

| Species | Embryonic mutation rate $\mu_E$ (mut/site/generation) | Mutation rate $\mu_O + \mu_S$ in the gametes after puberty (mut/site/year) | Age of puberty/ first reproduction (years) | Generation time $g$ (years) | Mutation rate $\mu = \mu_E + g \cdot (\mu_O + \mu_S)$ (mut/site/generation) |
| --- | --- | --- | --- | --- | --- |
| Human (38) | 6.26e-9 (95% C.I. 5.47e-9, 12.13e-9) | 3.5e-10 (95% C.I. 3.3e-10, 3.7e-10) | 13 | 30 | 1.2e-8 |
| Chimpanzee (39) | 5.11e-9 | 6.25e-10 | 14 | 25 | 1.2e-8 |
| Olive baboon (41) | 5.0e-9 | 1.4e-10 | 5.4 | 10 | 5.6e-9 |
| Rhesus macaque (42) | 3.9e-9 | 4.3e-10 | 3.5 | 8 | 5.8e-9 |
| Owl monkey (40) | 4.4e-9 | 6.6e-10 | 1 | 6.6 | 8.1e-9 |
| Domestic dog (45) | 3.75e-9 | 4.5e-10 | 1 | 4 | 5.1e-9 |
| Domestic cat (44) | 5.9e-9 | 8.2e-10 | 0.5 | 3.8 | 8.6e-9 |
| Mouse (48) | 3.75e-9 (95% C.I. 2.89e-9, 4.6e-9) | 1.64e-9 (95% C.I. 4.10e-10, 2.85e-9) | 0.15 | 0.75 | 4.7e-9 |

We performed log-log-linear regressions of *μ*_*E*_, *μ*_*O*_ + *μ*_*S*_, and *μ* = *μ*_*E*_ + *g* · (*μ*_*O*_ + *μ*_*S*_) as functions of generation time (log-log linear regressions are more appropriate than natural scale regressions because the distributions of generation times and mutation rate estimates are closer to lognormal than normal, as shown in **Supplementary Figure 1**). The regression results demonstrate that *μ*_*E*_ is positively correlated with generation time across these species, though less correlated with generation time than the raw germline mutation rate *μ* (**Figure 2A**). In contrast, the gamete mutation rate *μ*_*O*_ + *μ*_*S*_ is inversely correlated with generation time (**Figure 2B**). We performed all three of these regressions using a phylogenetic least squares (PGLS) approach but found that these traits had no phylogenetic signal across this small dataset (Pagel’s *λ* = 0), indicating that standard linear regression is also appropriate (see **Supplementary Table 1** for details). This result echoes recent findings of inverse correlations between lifespan and somatic mutation rates, a pattern that is hypothesized to result from selective pressure to moderate cancer risk and age-related decline in long-lived species (49– 51).

**Figure 2:**
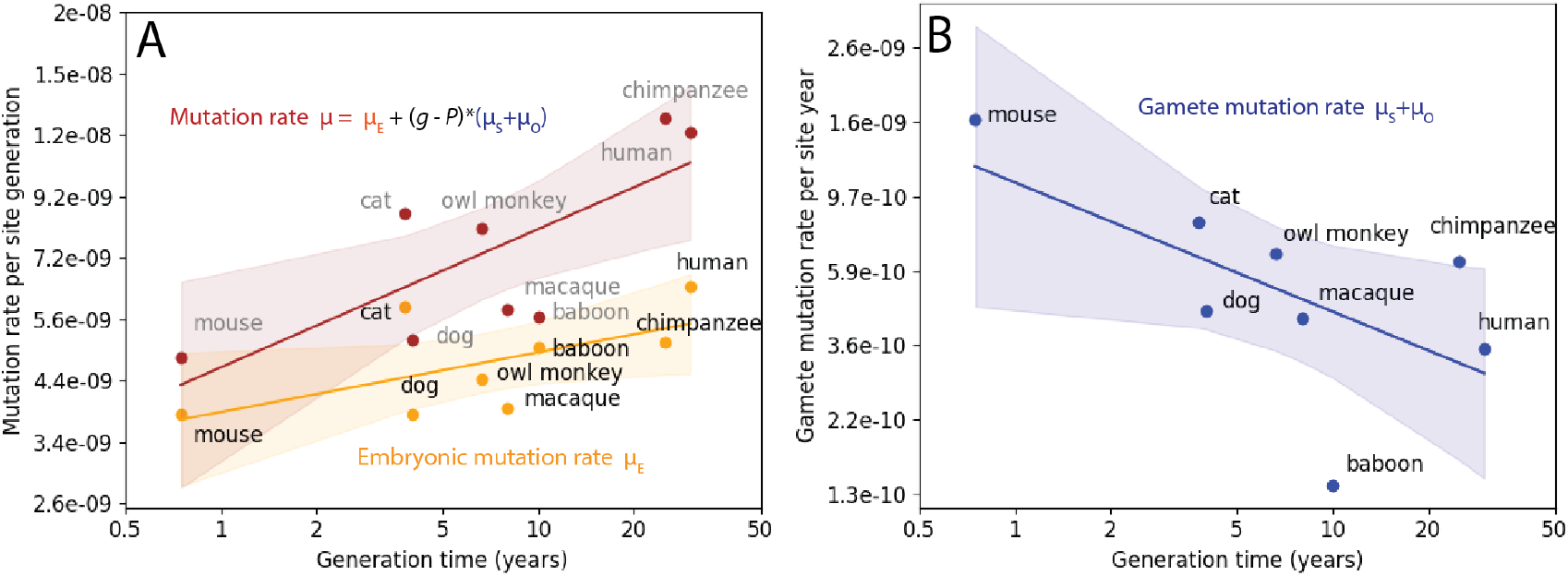
Variation among mammals in the rates of germline mutations occurring in the embryo and the gametes. **A**. Both the early embryonic mutation rate *μ*_*E*_ and the total mutation rate *μ* = *μ*_*E*_ + *g* · (*μ*_*O*_ + *μ*_*S*_) are positively correlated with the generation time *g* as measured by log-log linear regression. **B**. The mutation rate per year in the spermatocytes and oocytes post-puberty, *μ*_*O*_ + *μ*_*S*_, is negatively correlated with generation time as measured by a log-log linear regression.

### A “relaxed clock” reproductive longevity model predicts mutation rate variation across the full range of vertebrate lifespans

Figure 2. suggests that *μ*_*E*_ and *μ*_*O*_ + *μ*_*S*_ are not invariant among vertebrate species, but instead depend on generation time due to factors such as cell division rates, environmental mutagens, or the molecular efficacy of DNA repair. That being said, **Figure 2** contains data from only a handful of species due to the limited availability of suitable data for directly estimating *μ*_*E*_ and *μ*_*O*_ + *μ*_*S*_. Estimates of the overall germline mutation rate *μ* are available for many more species, and we hypothesized that the relationship among generation time, *μ*_*E*_, and *μ*_*O*_ + *μ*_*S*_ might translate into some constraints on the overall relationship between *g* and *μ*. Motivated by this, we developed a test to evaluate the fit of an empirical mutation rate distribution to either a strict, fixed-rate reproductive longevity model or a “relaxed clock” reproductive longevity model where *μ*_*E*_ and *μ*_*O*_ + *μ*_*S*_ are allowed to vary among species.

To formulate this test, we first approximated Equation (1) as a simple linear function of parental age by studying the relationship between age at first reproduction (a proxy for the timing of puberty) and average age at reproduction (a proxy for the generation time *g*) in a large set of vertebrate demographic data (52). In the notation of Equation (1), *g* equals both the paternal age *A*_*P*_ and the maternal age *A*_*M*_. We performed a linear regression of the age at first reproduction (*P*) against the average age at reproduction (*g*) and found that *P* is approximately equal to 0.42\**g* across 230 species with generation times ranging from 2 to 52 years (*r* = 0.87, see **Supplementary Figure 2**). Motivated by this, we further approximated Equation (1) using the assumption that *p* = *P/g* is a constant across species such that *g* − *P* = *g* · (1 − *P*/*g*) = *g* · (1 − *p*) and

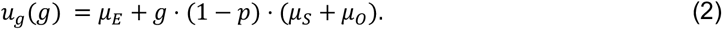

Letting *μ*_*E*_^*H*^, *μ*_*S*_^*H*^ and *μ*_*O*_^*H*^ be values of the embryonic, spermatocytic, and oocytic mutation rates estimated from human data, we substituted these values into (2) to predict mutation rate in the context of a strict reproductive longevity model that predicts the germline mutation rate *u*_*E*_ as a function of mutation rate parameters *μ*_*E*_^*H*^ and *μ*_*O*_^*H*^ + *μ*_*S*_^*H*^:

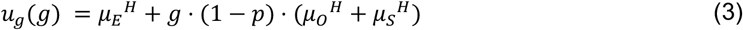

We then adapted equation (3) to formulate a relaxed-clock reproductive longevity model that allows the rates *μ*_*E*_ and *μ*_*O*_ + *μ*_*S*_ to vary as inferred from our meta-analysis in **Figure 2**. To capture variation in *u*_*E*_ as a function of generation time *g*, we let *u*_*E*_^(*g*)^ denote the early embryonic mutation rate at a generation time of *g* and let *α* denote the slope relating *log μ*_*E*_^(*g*)^ to *log g*. By these definitions,

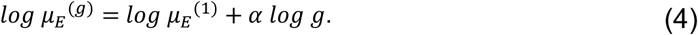

Exponentiating both sides of Equation (4) yields:

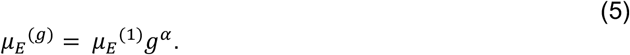

To capture gamete mutation rate variation in a similar way, we let *β* denote the slope of the regression relating *log* (*μ* _*S*_^(*g*)^ + *μ* _*O*_ ^(*g*)^) to *log g*, such that

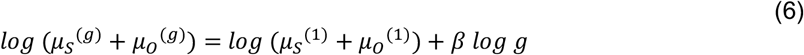

and

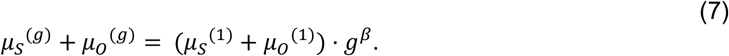

Substituting these values into equation (3) yields a prediction of the overall mutation rate:

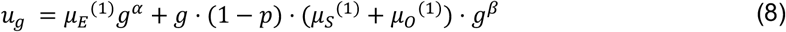

This simplifies to

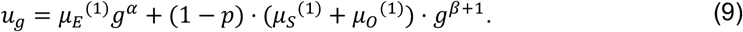

Equations (3) and (9) make two different concrete predictions about how mutation rates should vary with generation time among vertebrates. We were able to compare the accuracy of these predictions using a large vertebrate mutation rate dataset that was recently compiled by Wang and Obbard (26). As shown in **Figure 3**, the mutation rate per generation curve predicted by Equation (9) closely approximates the PGLS correlation between mutation rate and generation time. In contrast, the human constant-rate reproductive longevity model (Equation (3) with human-trained parameters) overestimates the mutation rates of species with short generation times. We also substituted mouse mutation rate parameters into (3) and found that the resulting model fits the mutation rates of short-generation-time vertebrates but overestimates the mutation rates of species with longer generation times. Both the human and mouse reproductive longevity models have greater upward concavity than the relaxed clock model: these models predict a relatively constant mutation rate for generation times less than 1 year, which is the generation time range where these models predict that almost all germline mutations occur in the embryo rather than the gametes.

**Figure 3:**
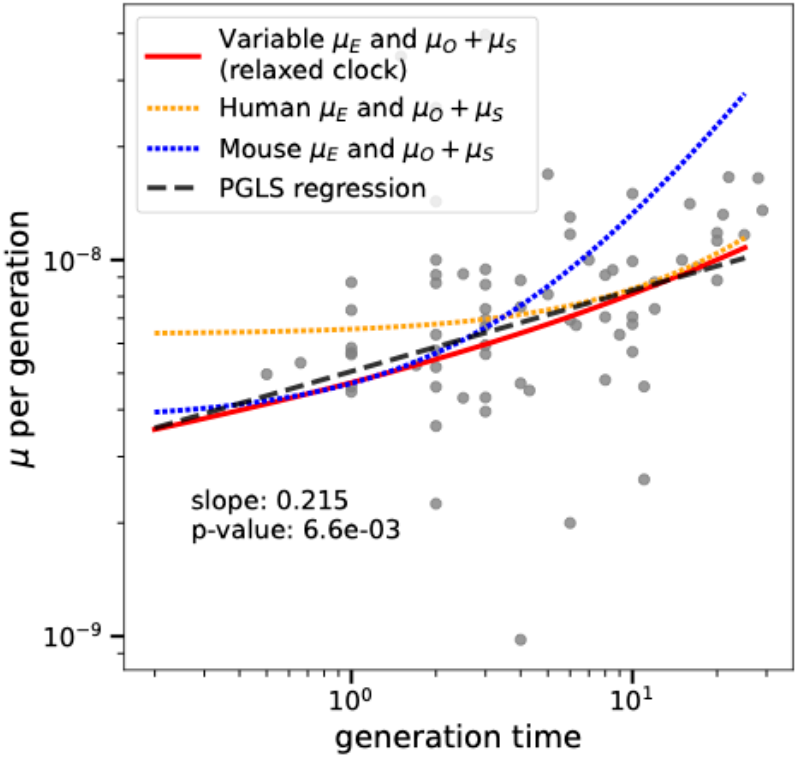
The relaxed-clock reproductive longevity model explains the correlation between mutation rate and generation time. A dashed line shows the PGLS regression of mutation rate versus generation time in vertebrates from Wang and Obbard’s mutation rate meta-analysis (26). This is close to the prediction of the relaxed rate reproductive longevity model fit to the multispecies pedigree data (solid red line). The prediction of the fixed-rate reproductive longevity model with human parameters (orange dotted line) overestimates the mutation rates associated with short generation times, while the fixed-rate reproductive longevity model with mouse parameters (blue dotted line) overestimates mutation rates associated with long generation times.

### Long lifespan increases the efficacy of selection for a low mutation rates in the gametes as well as the soma

So far, we have shown that vertebrate mutation rate variation is well described by a relaxed-clock reproductive longevity model where the early embryonic mutation rate per generation increases with reproductive age and the mutation rate per year in the gametes decreases with reproductive age. We will now go on to show that both the gamete mutation rate *μ*_*O*_ + *μ*_*S*_ and the embryonic mutation rate *μ*_*E*_ appear to be evolving in accord with the predictions of the drift-barrier hypothesis, with appropriate modification.

The drift barrier hypothesis explains the inverse correlation between mutation rate and *N*_*e*_ as a consequence of selection against weakly deleterious mutator alleles (19, 53). Mutator alleles might directly perturb DNA repair or proofreading, or they might indirectly affect the mutation rate by perturbing a trait like metabolism. Species with larger effective population sizes are generally better able to eliminate weakly deleterious alleles, while species with small effective sizes are more likely to retain these alleles as a result of stronger genetic drift (54). This leads to the prediction that mutator alleles will be more prevalent in low-*N*_*e*_ species, which also tend to have long generation times (55, 56). The gamete mutation rate *μ*_*O*_ + *μ*_*S*_ seems to contradict this prediction: we can extrapolate from **Figure 2B** that species with the smallest effective population sizes are somehow the most effective at eliminating gamete mutator alleles. We can explain this contradiction by looking more closely at how the fitness effect of a mutator allele is calculated.

Let *S*_*u*_^(*g*)^ be the selection coefficient of a mutator allele that creates *u* additional mutations per generation. Lynch previously estimated *S*_*u*_^(*g*)^ as follows (57): if *L* is the length of the diploid genome and each mutation has an expected fitness cost of E[*s*], then the expected selective cost of the mutator allele each generation is

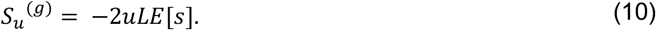

In a population of effective size *N*_2_, selection is predicted to eliminate mutations for which |*S*_*u*_^(*g*)^| > 1/(2*N*_2_). By this logic, natural selection should eliminate mutators whose per-generation mutation load *u* mutations per genome per generation satisfies the inequality

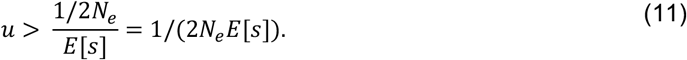

If we assume that *u, N*_*e*_, *L*, and *E*[*s*] are essentially independent variables, then as *N*_*e*_ gets larger, it will get progressively more difficult for a mutator to satisfy inequality (11) and thus the population should get more effective at purging away mutator alleles. A caveat is that this argument does not account for statistical dependence among *u, N*_*e*_, and the generation time *g*. We can reasonably assume that *u* and *g* are independent when considering a mutator allele that modifies *μ*_*E*_, since such a mutator will create the same embryonic mutation load regardless of when parents reproduce. However, for a mutator allele that alters *μ*_*S*_ + *μ*_*O*_ by creating extra mutations during spermatogenesis or oogenesis, the total mutation load created by the mutator each generation will scale proportional to *g*, as illustrated in **Figure 4A**. This will shift the distribution of mutator allele fitness effects toward more deleterious values in species with long generation times, an idea that Lindsay et al. previously posited to explain why mice have higher per-year germline mutation rates than humans do (48). We will refer to such a modifier of *μ*_*S*_ + *μ*_*O*_ as a “clocklike” mutator, in contrast to a “non-clocklike” mutator that modifies *μ*_*E*_ by a fixed amount each generation.

**Figure 4:**
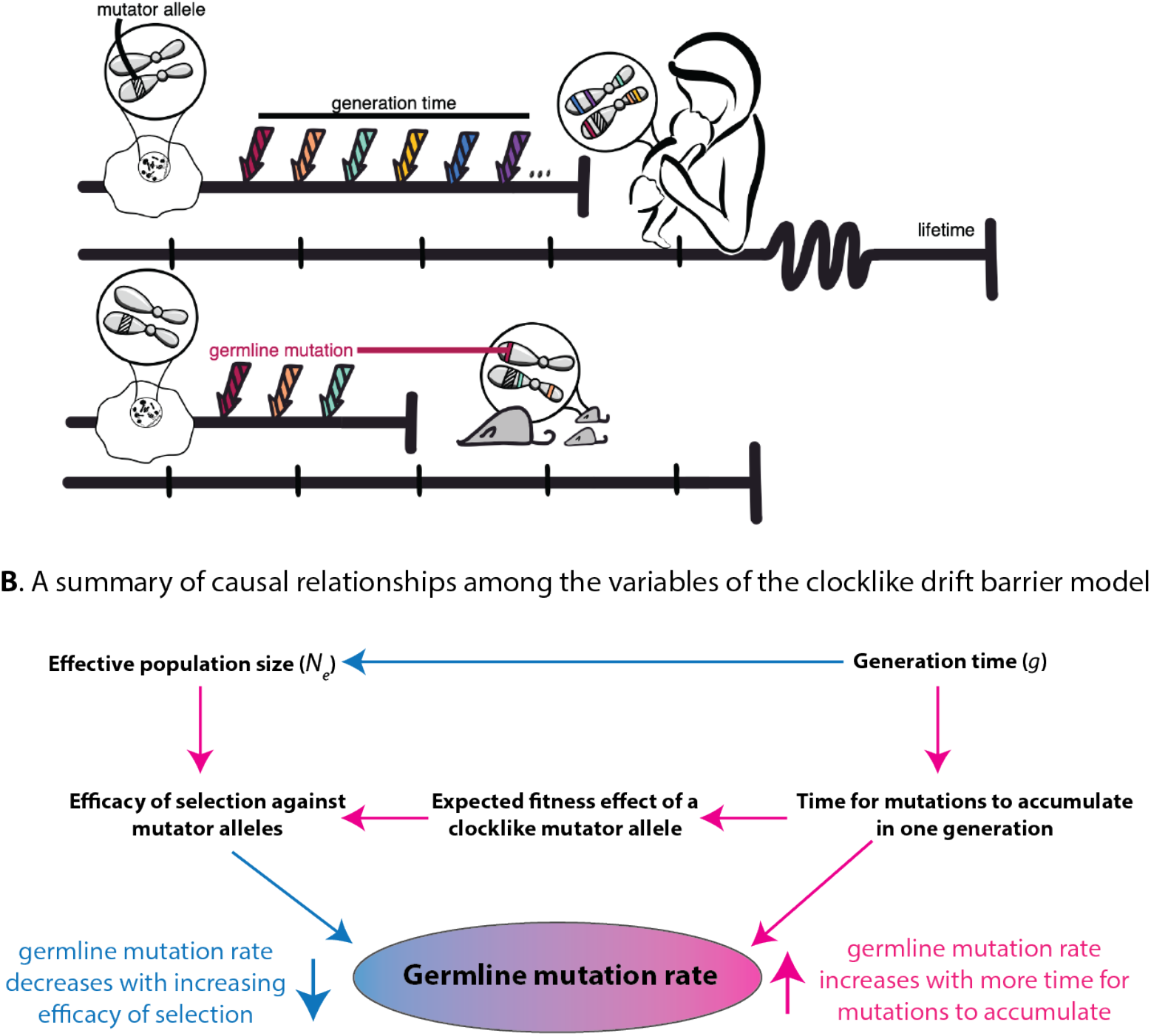
A model of germline mutation rate variation as a function of generation time, effective population size, and genetic variation that impacts the mutation rate measured per year. **A**. Here, we compare the effects of identical molecular changes occuring in some human DNA repair gene as well as its mouse homolog. If these mutator alleles produce the same number of germline mutations per year, the human allele will produce a greater mutation burden per generation compared to the mouse allele, leading to a greater expected fitness cost and a larger negative selection coefficient in the long-generation-time species. *Figure credit: Natalie Telis*. **B**. This diagram summarizes the multiple ways that generation time can affect the mutation rate, including its direct impact on the number of mutations that accumulate in a generation and its other impacts on the effective population size and the efficiency of natural selection. Pink arrows indicate positive correlations (an increase in the upstream variable causes an increase in the downstream variable), and blue arrows indicate negative correlations (an increase in the upstream variable causes a decrease in the downstream variable).

For a clocklike mutator that creates *k* additional mutations per year after puberty, the total fitness impact *S*_*k*_^(*y*)^ per generation will be the proportional to *k* times the number of years that elapse between puberty and reproduction in a generation of length *g*, which is *g*(1 − *p*). If the average fitness cost of a single mutation is *E*[*s*], then the total fitness impact of the mutator each generation will be

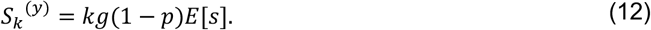

Since *S*_*k*_^(*y*)^ is proportional to the generation time *g*, this implies that as generation time increases, selection against clocklike mutators may get stronger, decreasing the mutation rate per year in the gametes and explaining the trend in **Figure 2B**. In order for the clocklike mutator to persist in the population, it must satisfy the familiar inequality *S*_*k*_^(*y*)^ > 1/(2*N*_2_), which will only hold if

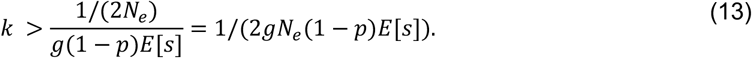

Inequality (11) defines a threshold of near-neutrality for modifiers of *μ*_*E*_, while (13) defines a threshold of near-neutrality for modifiers of *μ*_*S*_ + *μ*_*O*_ . If we ignore *E*[*s*] and *p*, assuming that these parameters do not vary much among species, then we conclude that the efficacy of selection against modifiers of *μ*_*E*_ is determined by *N*_*e*_ alone, while the efficacy of selection against modifiers of *μ*_*S*_ + *μ*_*O*_ is determined by the product *gN*_*e*_. **Figure 4B** summarizes how *g* and *N*_*e*_ interact to shape the gamete mutation load.

Our calculations suggest that the species with the lowest gamete mutation rates with be the species for which *gN*_*e*_ is the largest. However, the inverse correlation between *g* and *N*_*e*_ means that it is not obvious which life history strategies will maximize *gN*_*e*_. To gain clarity, we note that the relationship between *N*_*e*_ and *g* was previously studied in some detail during the initial development of the nearly neutral theory, since it was needed to explain the consistency in molecular substitution rates across the tree of life (58, 59). In this context, Chao and Carr previously measured an inverse log-linear correlation between *N*_*e*_ and *g* (55). We were able to reproduce this log-linear relationship in the Wang and Obbard mutation rate data (26)(**Figure 5A**).

**Figure 5:**
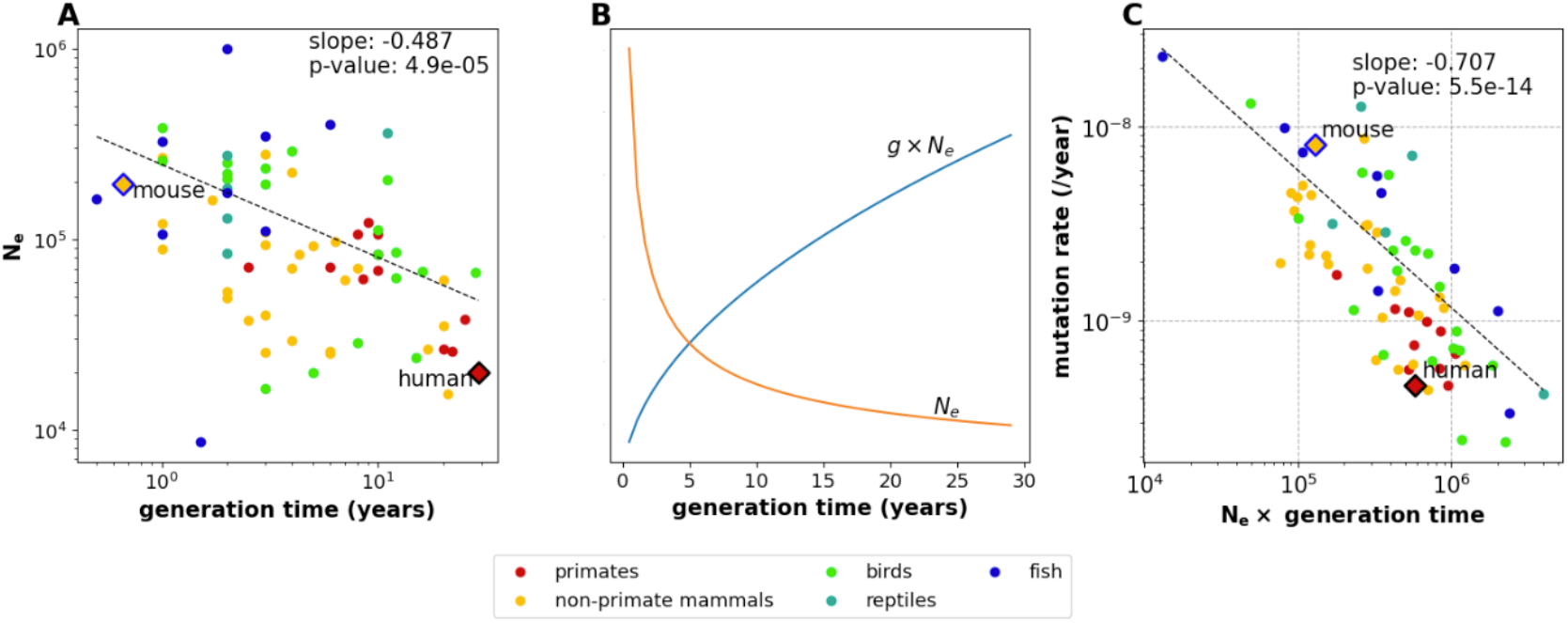
The relationship among *N*_*e*_, generation time, and the strength of selection against clocklike mutator alleles. **A**. The parameters log(*N*_*e*_) and log(*g*) are inversely correlated in the Wang and Obbard mutation rate data (26). We estimate a slope of -0.487 based on a PGLS regression. **B**. Expected values of *N*_*e*_ and *gN*_*e*_ as functions of *g*, extrapolated from the regression line in panel A and converted from log scale to natural scale. Each curve has been visualized using an arbitrary *y*-axis scaling, and together they illustrate that *gN*_*e*_ increases with increasing *g* even as *N*_*e*_ decreases. **C**. Mutation rate estimates from Bergeron, et al. confirm that the mutation rate per year decreases as a function of *gN*_*e*_, as expected if long generation times dominate the effect of decreasing effective population size to strengthen selection against clocklike mutator alleles. Note that the long-lived primates have higher values of *gN*_*e*_ than the short-lived, high-*N*_*e*_ mouse.

The linear relationship *log N*_*e*_ = *γ log g* +*log C* (where *γ* and *C* are constants) implies that *N*_*e*_ = *Cg*^*γ*^ and *gN*_*e*_ = *Cg*^1+*γ*^. This expression might increase or decrease with increasing *g* depending whether *γ* is greater or less than -1, so knowing the value of *γ* is key to deciding whether species with long or short generation times are likely to have the lowest mutation rates. We estimate that *γ* ≈ −0.487 based on a PGLS regression of log(*N*_*e*_) against log(*g*), remarkably close to the value of -0.5 that Kimura and Ohta originally proposed to reconcile the nearly neutral theory with the molecular clock model (55, 60). This implies that g*gN*_*e*_ = *g*^1+*γ*^ = *g*^1−0.487^ = *g*^0.513^. As shown in **Figure 5B**, this implies that *gN*_*e*_ behaves approximately like 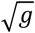, increasing as *g* increases. Therefore, if we compare fast-reproducing species like mice to slower-reproducing species like humans, the slower-reproducing species will have smaller values of *N*_*e*_ but larger values of *gN*_*e*_, which is the parameter that determines the strength of selection for a low mutation rate per year in the gametes. **Figure 5C** shows empirically that *gN*_*e*_ is negatively correlated with the germline mutation rate per year, consistent with the idea that the parameter *gN*_*e*_ determines the strength of selection against mutator alleles. We can also see that humans and other long-lived primates have high values of *gN*_*e*_ compared to the short-lived mouse.

## Discussion

We have introduced a framework for combining two models of mutation rate evolution, the reproductive longevity model and the drift-barrier model, into a relaxed-clock reproductive longevity model that explains the nuanced relationship between mutation rate and reproductive age. The early embryonic mutation rate appears to have been pushed to its lowest levels in species with the largest effective population sizes, consistent with the predictions of the nearly neutral theory. In contrast, the gamete mutation rate trends in the opposite direction, achieving its lowest levels in long-lived animals with small effective population sizes. This is consistent with our argument that long generation times should intensify the strength of selection against clocklike mutator alleles, overcoming the tendency of small effective population sizes to dampen the general effectiveness of selection.

Variation in the gamete mutation rate per year appears to echo patterns of mutation rate variation in somatic tissues. A recent study of colon crypt mutations found an inverse log-log linear relationship between lifespan and the mutation rate per year (51), mirroring the correlation we observe between generation time and the mutation rate in the gametes. In both cases, the fitness effect of any mutation rate increase becomes compounded over the lifetime of the cell lineage that is mutating, giving long-lived, late-reproducing organisms a stronger incentive to preserve genomic integrity (61, 62). In gerontology, this concept is known as the disposable soma theory (63, 64), and our analysis suggests that a version of this theory is also applicable to renewing germline tissues. Since the same molecular machinery is ultimately responsible for safeguarding both germline and somatic DNA, pleiotropy between somatic and germline mutation rates may amplify differences among species in the strength of selection against clocklike mutator alleles.

While selection against nearly neutral mutator alleles is a parsimonious explanation for the observation that longer generation times are associated with higher rates of embryonic mutations and lower rates of gamete mutations, other explanations are also possible. Later reproduction is generally associated with a larger body size and longer gestation, either of which might cause additional mutations to accumulate in the embryonic germline. It is also possible that the higher gamete mutation rate in fast-reproducing organisms might be driven by biological factors such as higher metabolism or higher sperm production volume. These alternate hypotheses may become testable as additional generation-time-calibrated mutation rate estimates become available. Our theoretical work underscores the value of collecting mutation rate data in a way that facilitates separate estimation of embryonic and germ cell mutation rates, whether by sequencing multi-offspring pedigrees (65–67) or using emerging technologies such as single-cell gamete sequencing (68, 69).

Recent research on de novo mutagenesis has built a multifactorial case that most mutations are products of DNA damage rather than cell division error (41, 47, 70–72). However, embryonic mutations might be the exception to this rule if they largely originate during a few error-prone postzygotic cell divisions. Human and mouse DNM data, which are higher resolution than the data available for any other species, make it clear that early embryonic cell divisions have elevated mutation rates (34, 73–76), possibly due to the reliance of this early-stage embryo on maternal DNA repair prior to the maternal-zygotic transition (73, 74). However, Drost and Lee have argued that most mammals, including mice and humans, have similar primordial germ cell developmental trajectories, with similar numbers of cell divisions leading from the zygote to the germ cells (32). This implies that variation in the rate of embryonic mutations among mammals is not likely driven by variation in the number of early embryonic cell divisions but is more likely driven by variation in DNA damage or repair during early development. Primordial germ cell specification occurs around gastrulation, which takes place between 6 and 9 days of embryonic development in mouse (77) and between 14 and 21 days of embryonic development in humans (78). It is possible that the slower pace of early development in longer-lived vertebrates allows more unrepaired DNA damage to accumulate and drives the tendency of longer-lived vertebrates to have higher rates of early embryonic mutations.

In addition to making testable predictions about the molecular efficacy of DNA repair and how it varies among species, our model provides a straightforward way to impute the germline mutation rates of species for which direct measurements are missing. If a species’ age of reproductive maturity and average generation time have both been estimated, Equation (9) provides a mutation rate estimate that can be used for calibrating phylogenetic trees and demographic histories. Although such a mutation rate estimate will not be as accurate as a mutation rate estimated directly from trio sequencing data, it may be more reliable than attempting to infer the mutation rate from phylogenetic data, which famously overestimated the human mutation rate by a factor of 2 (79–81) and also reached inaccurate conclusions about baleen whale mutation rates (82). Our model may even be useful for imputing the mutation rates of non-mammalian species; for example, the mutation rate of the black abalone is similar to the mutation rates of vertebrates with similar reproductive lifespans (83). We have not attempted here to deduce how mutation rates are affected by body size (84), domestication history (85), or the countless other variables that may affect genomic integrity, but a good model encapsulating the effects of generation time should improve our power to learn the effects of additional variables in the future.

## Methods

### Meta-analysis of mutation rates from mammalian pedigrees

We obtained estimates of the embryonic mutation rate (*μ*_*E*_) and the gamete mutation rate per year after puberty (*μ*_*O*_ + *μ*_*S*_) from eight mammalian pedigree studies. Each study performed a regression of mutation rate against paternal and/or maternal age, but the studies reported the regression results in a variety of different ways. Below we report how each study’s age regression parameters were transformed into estimates of *μ*_*E*_ and *μ*_*O*_ + *μ*_*S*_.

#### Human

Our human mutation parameter estimates are derived from Jonsson, et al. 2017 (38), Supplementary Table 6, which gives the maternal slope *m*_*s*_, maternal intercept *m*_*i*_, paternal slope *p*_*s*_, and paternal intercept *p*_*i*_ of the paper’s Poisson regression of the dependence of mutation rate on parental age (maternal and paternal intercepts represent the interpolated maternal and paternal mutation loads at a reproductive age of zero years). Upper and lower 95% confidence bounds for each of these variables are also given. The accessible haploid genome size *A* is listed as 2722501677 base pairs in the caption of Supplementary Table 17. We calculated *μ*_*E*_, the mutation load at puberty (age 13) and *μ*_*O*_ + *μ*_*S*_, the mutation rate per year in the gametes post puberty, as follows:

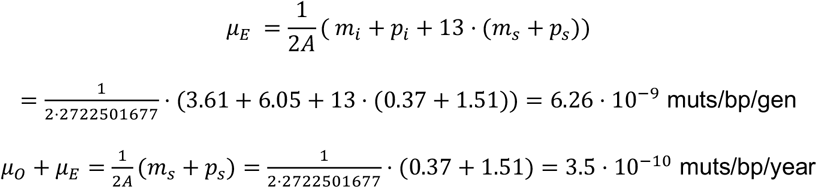

The upper and lower confidence bounds on *μ*_*E*_ and *μ*_*O*_ + *μ*_*S*_ were calculated in the same way using the upper and lower bounds of the regression parameters.

#### Chimpanzee

Venn, et al. (39) reported a chimpanzee paternal age effect of 2.95 additional mutations per site per year and a maternal age effect of zero additional mutations per site per year (all regression parameter estimates are given in Table S10). They reported a paternal intercept of -23.8 total mutations per generation and a flat maternal contribution of 6.65 mutations per generation. The earliest reproductive age reported in the data is 14 years, and the size of the accessible haploid genome is reported to be 2360 megabases. Using these parameters, we calculated that:

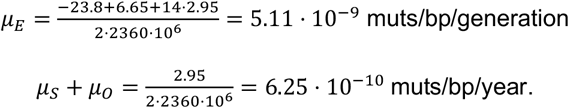

#### Olive baboon

Wu, et al. 2020 (41) reported a paternal slope of 0.15 DNMs per genome per year and a maternal slope of 0.65 DNMs per genome per year (see results section “Estimating sex-specific germline mutation rates and age effects”). These values are scaled to a haploid genome size of 2.581 · 10^9^ base pairs, from which we calculate that

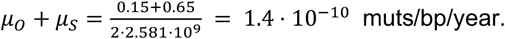

To calculate *μ*_*E*_, we used the regression coefficients reported in S2 Data, Fig 2B. The reported maternal intercept is 0.23 mutations per genome at a maternal age of 0.55 years, and the reported paternal intercept is 22.16 mutations per genome at a paternal age of 0.15 years. Supplementary Table 14 reports an age of male puberty of 5.41 years, so we estimated the mutation load at puberty by adding the maternal and maternal intercepts to the estimated maternal and paternal mutation load accumulated in a period of 5 years. Dividing this load by the genome size, we obtain:

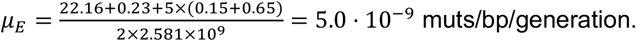

#### Rhesus macaque

Wang, et al. 2020 (42) report a total parental age slope of 4.3 · 10^−10^ mutations per site per year and a mutation load at puberty of 3.9 · 10^−9^ mutations per site per generation. We were able to use these values without further transformation. A second linear model of macaque mutation rate as a function of generation time was generated by Bergeron, et al. (43), but we chose to use the Wang et al. model for consistency with the pipeline that was used to generate the owl monkey and domestic cat mutation rate models.

#### Owl monkey

Equation (2) in Thomas, et al. 2018 (40) reports a parental age slope of *μ*_*O*_ + *μ*_*S*_ = 6.62 · 10^−10^ mutations per site per year and y-intercept of 3.74 · 10^−9^. We estimate a pre-puberty mutation load *μ*_*E*_ = 4.40 · 10^−9^ assuming a generation time of 1 year and adding a year of gamete mutation accumulation to the *y*-intercept. Since Thomas, et al. report paternal and maternal generation times of 6.64 and 6.53, we use an owl monkey generation time of 6.6 years.

#### Domestic cat

Wang et al. 2022 (44) report mutation rates of *μ*_*E*_ = 5.9 × 10^−9^ per site per generation for reproduction at the age of puberty and an overall average mutation rate of 8.6 × 10^−9^ mutations per site per generation. They assume that puberty occurs at 0.5 years and report an average reproductive age of 3.8 years in their data. Using these values we calculate that

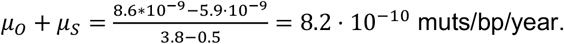

#### Domestic dog

Figure 2b of Zhang, et al. 2024 (45) shows bar plot representations of the slopes and intercepts defining the maternal and paternal mutation rates as linear functions of reproductive age. Since numerical estimates of these parameters are not reported in the text, we extrapolated them from the bar plot heights. The maternal mutation rate slope and intercept appear to be 1 × 10^−10^ and 8 × 10^−10^, while the paternal mutation rate slope and intercept appear to be 3.5 × 10^−10^ and 2.5 × 10^−9^. Assuming an age of 1 year at puberty (which appears to be the minimum age at first reproduction represented in the dataset) we conclude that:

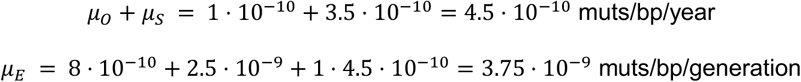

#### Mouse

We downloaded the supplementary mutation data from Lindsay, et al. 2019 (48), which reports accessible-genome-corrected mutation counts and parental age at conception in weeks for all of the offspring in their pedigrees. We performed a regression of mutation rate against parental age and used the results to calculate means and confidence intervals for murine *μ*_*O*_ + *μ*_*S*_ and *μ*_*E*_.

### Meta-analysis of the correlation between mutation rate per year and generation time

We used the nucleotide diversity (*π*),mutation rate (*μ*) and generation time (*g*) data compiled by Wang and Obbard to quantify the relationship between *g* and *N*_*e*_. We first estimated *N*_*e*_ for each species via the formula *N*_*e*_ = *π*/94 · *μ*8 (26) (see Data and Code Availability). We then performed a PGLS regression of mutation rate against *g* · *N*_*e*_ using the R library caper (86). Additionally, we estimated Pagel’s *λ* (87) to be 0.92 using caper’s maximum likelihood implementation. *λ* is commonly used to quantify the amount of phylogenetic signal in the dataset. It is a scaling parameter applied to internal branch lengths in the phylogenetic tree, and is typically a value between 0 and 1. *λ* = 1 means that the traits being regressed against one another appear to have evolved according to a Brownian motion evolutionary model and is interpreted as strong evidence for phylogenetic signal in the dataset, whereas *λ* = 0 suggests that the traits evolved completely independently of the phylogenetic tree structure. See **Supplementary Table 1** for detailed numerical regression results.

## Competing Interest Statement

The authors declare no competing interests.

## Data and Code Availability

The mutation rates, nucleotide diversity, generation time data, and phylogenetic tree utilized in our calculations were originally compiled by Wang and Obbard (26) and are all publicly available at https://github.com/Yiguan/mutation_literature. The code we used to perform this paper’s analysis is available at https://github.com/harrispopgen/clocklike-DBH.

## Acknowledgements

We thank Joshua Schraiber for manuscript comments, and we thank Michael Lynch and members of the Harris Lab for helpful advice and discussions. We also thank Natalie Telis for figure design assistance. K.H. received support from NIH/NIGMS grant R35GM133428, a Burroughs Wellcome Fund Career Award at the Scientific Interface, a Searle scholarship, a Pew Scholarship, the Allen Discovery Center for Cell Lineage Tracing, and a Sloan Fellowship. A.C.B. received additional support from the National Institute of Health (NIH) “Biological Mechanisms of Healthy Aging” training program (T32 AG066574).

## Supplementary Figures

**Supplementary Figure 1:**
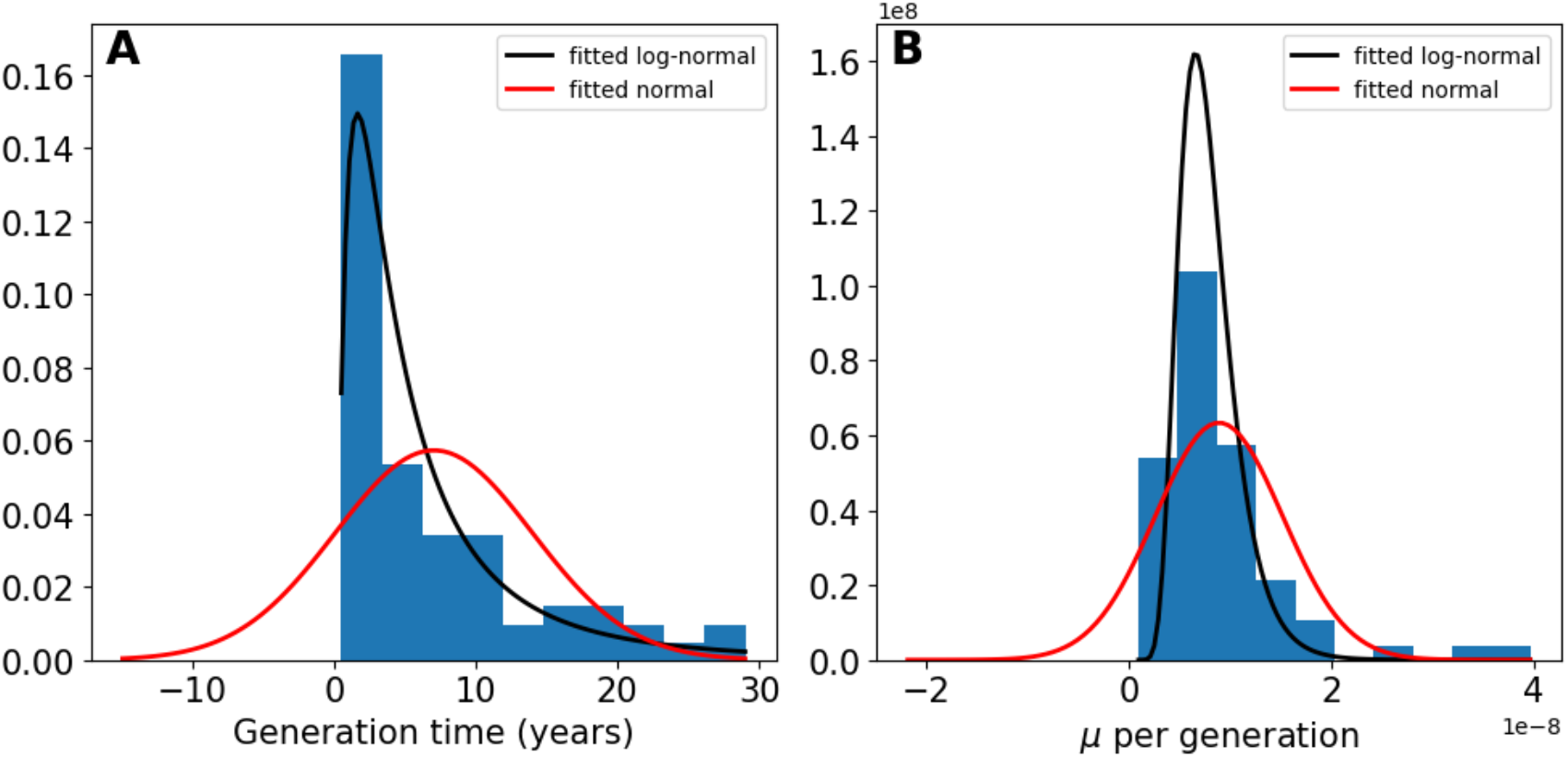
Distributions of generation time and mutation rate per generation across species. Data taken from Wang and Obbard (26). Red and black lines correspond to the fitted normal and log-normal distributions, respectively. Lognormal provides a better fit to both the distribution of generation times and the distribution of the mutation rate per generation.

**Supplementary Figure 2:**
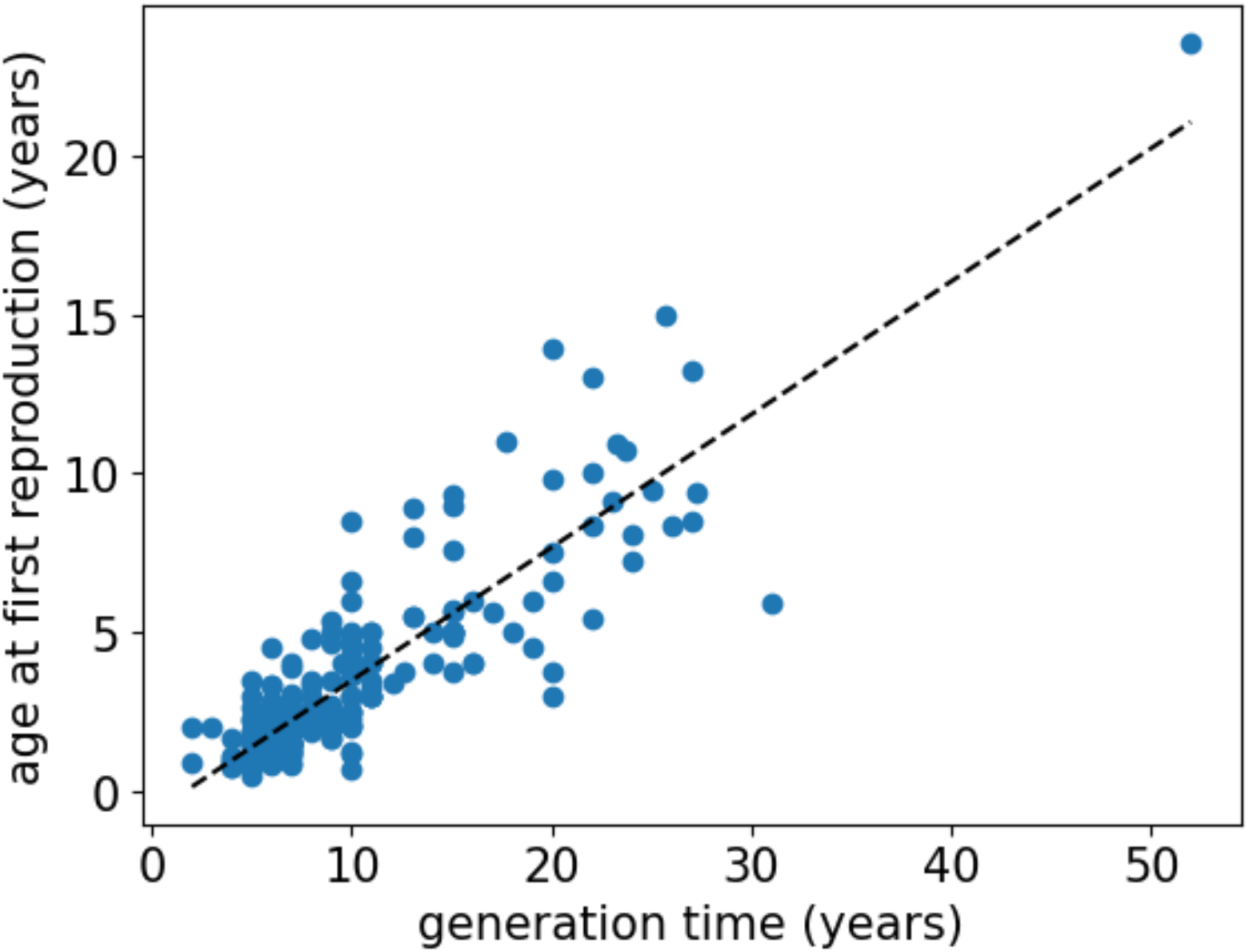
Regression of age at first reproduction versus generation time. Data taken from Pacifici et al. (52). Age at first reproduction is used as a proxy for age at puberty. Age at first reproduction is found to be linear with respect to generation time, with a slope of 0.42.

## Notes

### Competing Interest Statement

The authors have declared no competing interest.

### Summary of Updates

The manuscript has been extensively revised. While the original manuscript argued that the germline mutation rate as a whole evolves in a way that is dependent on generation time, the revised manuscript decomposes the mutation rate into embryonic and gamete mutation rates and then uses variation in parental age effects among species to show that gamete mutation rate evolve to be lower as generation time increases, while embryonic mutation rates show the opposite pattern. Figures 1, 2, and 3 are new figures describing this model framing, while Figures 4 and 5 are carried over from the original manuscript.

## References

1. E. G. Leigh, The evolution of mutation rates. Genetics 73, Suppl 73:1–18 (1973).

2. C. F. Baer, M. M. Miyamoto, D. R. Denver, Mutation rate variation in multicellular eukaryotes: causes and consequences. Nat. Rev. Genet. 8, 619–631 (2007).

3. J. J. Bull, R. Sanjuán, C. O. Wilke, Theory of Lethal Mutagenesis for Viruses. J. Virol. (2007). 10.1128/jvi.01624-06.

4. M. Eigen, P. Schuster, The Hypercycle: A Principle of Natural Self-Organization (Springer Science & Business Media, 2012).

5. M. Lynch, et al., Genetic drift, selection and the evolution of the mutation rate. Nat. Rev. Genet. 17, 704–714 (2016).

6. M. A. Bedau, N. H. Packard, Evolution of evolvability via adaptation of mutation rates. Biosystems 69, 143–162 (2003).

7. J. L. Payne, A. Wagner, The causes of evolvability and their evolution. Nat. Rev. Genet. 20, 24–38 (2019).

8. W. Wei, et al., Rapid evolution of mutation rate and spectrum in response to environmental and population-genetic challenges. Nat. Commun. 13, 4752 (2022).

9. M. Lynch, R. Bürger, D. Butcher, W. Gabriel, The Mutational Meltdown in Asexual Populations. J. Hered. 84, 339–344 (1993).

10. C. Zeyl, M. Mizesko, J. A. G. M. De Visser, Mutational Meltdown in Laboratory Yeast Populations. Evolution 55, 909–917 (2001).

11. J. W. Drake, A constant rate of spontaneous mutation in DNA-based microbes. Proc. Natl. Acad. Sci. 88, 7160–7164 (1991).

12. A. S. Kondrashov, Modifiers of mutation-selection balance: general approach and the evolution of mutation rates. Genet. Res. 66, 53–69 (1995).

13. P. D. Sniegowski, P. J. Gerrish, T. Johnson, A. Shaver, The evolution of mutation rates: separating causes from consequences. BioEssays 22, 1057–1066 (2000).

14. M. Lynch, G. K. Marinov, The bioenergetic costs of a gene. Proc. Natl. Acad. Sci. 112, 15690–15695 (2015).

15. L. Bromham, A. Rambaut, P. H. Harvey, Determinants of rate variation in mammalian DNA sequence evolution. J. Mol. Evol. 43, 610–621 (1996).

16. B. Nabholz, S. Glémin, N. Galtier, Strong Variations of Mitochondrial Mutation Rate across Mammals—the Longevity Hypothesis. Mol. Biol. Evol. 25, 120–130 (2008).

17. L. Bromham, The genome as a life-history character: why rate of molecular evolution varies between mammal species. Philos. Trans. R. Soc. B Biol. Sci. 366, 2503–2513 (2011).

18. M. Kimura, On the evolutionary adjustment of spontaneous mutation rates. Genet. Res. 9, 23–34 (1967).

19. W. Sung, M. S. Ackerman, S. F. Miller, T. G. Doak, M. Lynch, Drift-barrier hypothesis and mutation-rate evolution. Proc. Natl. Acad. Sci. 109, 18488–18492 (2012).

20. M. Lynch, et al., The divergence of mutation rates and spectra across the Tree of Life. EMBO Rep. 24, e57561 (2023).

21. B. H. Good, M. M. Desai, Evolution of Mutation Rates in Rapidly Adapting Asexual Populations. Genetics 204, 1249–1266 (2016).

22. W. R. Milligan, G. Amster, G. Sella, The impact of genetic modifiers on variation in germline mutation rates within and among human populations. Genetics 221, iyac087 (2022).

23. M. Lynch, The evolutionary scaling of cellular traits imposed by the drift barrier. Proc. Natl. Acad. Sci. 117, 10435–10444 (2020).

24. A. H. Sturtevant, Essays on Evolution. I. On the Effects of Selection on Mutation Rate. Q. Rev. Biol. 12, 464–467 (1937).

25. K. Harris, J. K. Pritchard, Rapid evolution of the human mutation spectrum. eLife 6, e24284 (2017).

26. Y. Wang, D. J. Obbard, Experimental estimates of germline mutation rate in eukaryotes: a phylogenetic meta-analysis. Evol. Lett. 7, 216–226 (2023).

27. L. A. Bergeron, et al., Evolution of the germline mutation rate across vertebrates. Nature 615, 285–291 (2023).

28. N. A. Moran, H. J. McLaughlin, R. Sorek, The Dynamics and Time Scale of Ongoing Genomic Erosion in Symbiotic Bacteria. Science 323, 379–382 (2009).

29. M. S. Snoke, T. U. Berendonk, D. Barth, M. Lynch, Large Global Effective Population Sizes in Paramecium. Mol. Biol. Evol. 23, 2474–2479 (2006).

30. W. Sung, et al., Extraordinary genome stability in the ciliate Paramecium tetraurelia. Proc. Natl. Acad. Sci. 109, 19339–19344 (2012).

31. H. Long, T. G. Doak, M. Lynch, Limited Mutation-Rate Variation Within the Paramecium aurelia Species Complex. G3 GenesGenomesGenetics 8, 2523–2526 (2018).

32. J. B. Drost, W. R. Lee, Biological basis of germline mutation: Comparisons of spontaneous germline mutation rates among drosophila, mouse, and human. Environ. Mol. Mutagen. 25, 48–64 (1995).

33. B. Arbeithuber, A. J. Betancourt, T. Ebner, I. Tiemann-Boege, Crossovers are associated with mutation and biased gene conversion at recombination hotspots. Proc. Natl. Acad. Sci. 112, 2109–2114 (2015).

34. Y. S. Ju, et al., Somatic mutations reveal asymmetric cellular dynamics in the early human embryo. Nature 543, 714–718 (2017).

35. J. J. Welch, O. R. Bininda-Emonds, L. Bromham, Correlates of substitution rate variation in mammalian protein-coding sequences. BMC Evol. Biol. 8, 53 (2008).

36. N. Risch, E. W. Reich, M. M. Wishnick, J. G. McCarthy, Spontaneous mutation and parental age in humans. Am. J. Hum. Genet. 41, 218 (1987).

37. A. Kong, et al., Rate of de novo mutations and the importance of father’s age to disease risk. Nature 488, 471–475 (2012).

38. H. Jónsson, et al., Parental influence on human germline de novo mutations in 1,548 trios from Iceland. Nature 549, 519–522 (2017).

39. O. Venn, et al., Strong male bias drives germline mutation in chimpanzees. Science 344, 1272–1275 (2014).

40. G. W. C. Thomas, et al., Reproductive longevity predicts mutation rates in primates. Curr. Biol. 28, 3193-3197.e5 (2018).

41. F. L. Wu, et al., A comparison of humans and baboons suggests germline mutation rates do not track cell divisions. PLOS Biol. 18, e3000838 (2020).

42. R. J. Wang, et al., Paternal age in rhesus macaques is positively associated with germline mutation accumulation but not with measures of offspring sociability. Genome Res. 30, 826–834 (2020).

43. L. A. Bergeron, et al., The germline mutational process in rhesus macaque and its implications for phylogenetic dating. GigaScience 10 (2021).

44. R. J. Wang, et al., De novo mutations in domestic cat are consistent with an effect of reproductive longevity on both the rate and spectrum of mutations. Mol. Biol. Evol. 39, msac147 (2022).

45. S.-J. Zhang, et al., Determinants of de novo mutations in extended pedigrees of 43 dog breeds. [Preprint] (2024). Available at: https://www.biorxiv.org/content/10.1101/2024.06.04.596747v1 [Accessed 7 November 2024].

46. G. Amster, G. Sella, Life history effects on the molecular clock of autosomes and sex chromosomes. Proc. Natl. Acad. Sci. 113, 1588–1593 (2016).

47. Z. Gao, M. J. Wyman, G. Sella, M. Przeworski, Interpreting the Dependence of Mutation Rates on Age and Time. PLOS Biol. 14, e1002355 (2016).

48. S. J. Lindsay, R. Rahbari, J. Kaplanis, T. Keane, M. E. Hurles, Similarities and differences in patterns of germline mutation between mice and humans. Nat. Commun. 10, 4053 (2019).

49. B. Milholland, et al., Differences between germline and somatic mutation rates in humans and mice. Nat. Commun. 8, 15183 (2017).

50. L. Zhang, et al., Maintenance of genome sequence integrity in long- and short-lived rodent species. Sci. Adv. 7, eabj3284 (2021).

51. A. Cagan, et al., Somatic mutation rates scale with lifespan across mammals. Nature 604, 517–524 (2022).

52. M. Pacifici, et al., Generation length for mammals. Nat. Conserv. 5, 89–94 (2013).

53. M. Lynch, Evolution of the mutation rate. Trends Genet. 26, 345–352 (2010).

54. T. Ohta, The nearly neutral theory of molecular evolution. Annu. Rev. Ecol. Syst. 23, 263– 286 (1992).

55. L. Chao, D. E. Carr, The molecular clock and the relationship between population size and generation time. Evolution 47, 688–690 (1993).

56. R. S. Waples, G. Luikart, J. R. Faulkner, D. A. Tallmon, Simple life-history traits explain key effective population size ratios across diverse taxa. Proc. R. Soc. B Biol. Sci. (2013). 10.1098/rspb.2013.1339.

57. M. Lynch, The cellular, developmental and population-genetic determinants of mutation-rate evolution. Genetics 180, 933–943 (2008).

58. T. Ohta, Very slightly deleterious mutations and the molecular clock. J. Mol. Evol. 26, 1–6 (1987).

59. T. Ohta, H. Tachida, Theoretical study of near neutrality. I. Heterozygosity and rate of mutant substitution. Genetics 126, 219–229 (1990).

60. M. Kimura, Model of effectively neutral mutations in which selective constraint is incorporated. Proc. Natl. Acad. Sci. 76, 3440–3444 (1979).

61. R. Peto, F. J. Roe, P. N. Lee, L. Levy, J. Clack, Cancer and ageing in mice and men. Br. J. Cancer 32, 411–426 (1975).

62. M. Tollis, A. M. Boddy, C. C. Maley, Peto’s Paradox: how has evolution solved the problem of cancer prevention? BMC Biol. 15, 60 (2017).

63. L. Hayflick, “Current theories of biological aging” in Biology of Aging and Development, G. J. Thorbecke, Ed. (Springer US, 1975), pp. 11–19.

64. T. B. L. Kirkwood, R. Holliday, J. Maynard Smith, R. Holliday, The evolution of ageing and longevity. Proc. R. Soc. Lond. B Biol. Sci. 205, 531–546 (1997).

65. R. Rahbari, et al., Timing, rates and spectra of human germline mutation. Nat. Genet. 48, 126–133 (2016).

66. T. A. Sasani, et al., Large, three-generation human families reveal post-zygotic mosaicism and variability in germline mutation accumulation. eLife 8, e46922 (2019).

67. D. Porubsky, et al., A familial, telomere-to-telomere reference for human de novo mutation and recombination from a four-generation pedigree. bioRxiv 2024.08.05.606142 (2024). 10.1101/2024.08.05.606142.

68. A. D. Bell, et al., Insights into variation in meiosis from 31,228 human sperm genomes. Nature 583, 259–264 (2020).

69. M. D. Neville, et al., Sperm sequencing reveals extensive positive selection in the male germline. [Preprint] (2024). Available at: https://www.medrxiv.org/content/10.1101/2024.10.30.24316414v1 [Accessed 7 November 2024].

70. W. S. W. Wong, et al., New observations on maternal age effect on germline de novo mutations. Nat. Commun. 7, 10486 (2016).

71. Z. Gao, et al., Overlooked roles of DNA damage and maternal age in generating human germline mutations. Proc. Natl. Acad. Sci. 116, 9491–9500 (2019).

72. N. Spisak, M. de Manuel, W. Milligan, G. Sella, M. Przeworski, The clock-like accumulation of germline and somatic mutations can arise from the interplay of DNA damage and repair. PLOS Biol. 22, e3002678 (2024).

73. M. A. Eckersley-Maslin, C. Alda-Catalinas, W. Reik, Dynamics of the epigenetic landscape during the maternal-to-zygotic transition. Nat. Rev. Mol. Cell Biol. 19, 436–450 (2018).

74. E. V. Khokhlova, Z. S. Fesenko, J. V. Sopova, E. I. Leonova, Features of DNA Repair in the Early Stages of Mammalian Embryonic Development. Genes 11, 1138 (2020).

75. H. Lee, et al., Characterization of early postzygotic somatic mutations through multi-organ analysis. J. Hum. Genet. 66, 777–784 (2021).

76. A. Uchimura, et al., Early embryonic mutations reveal dynamics of somatic and germ cell lineages in mice. Genome Res. 32, 945–955 (2022).

77. E. S. Bardot, A.-K. Hadjantonakis, Mouse gastrulation: Coordination of tissue patterning, specification and diversification of cell fate. Mech. Dev. 163, 103617 (2020).

78. R. C. V. Tyser, et al., Single-cell transcriptomic characterization of a gastrulating human embryo. Nature 600, 285–289 (2021).

79. A. Scally, R. Durbin, Revising the human mutation rate: implications for understanding human evolution. Nat. Rev. Genet. 13, 745–753 (2012).

80. A. Scally, Mutation rates and the evolution of germline structure. Philos. Trans. R. Soc. B Biol. Sci. 371, 20150137 (2016).

81. P. Moorjani, Z. Gao, M. Przeworski, Human Germline Mutation and the Erratic Evolutionary Clock. PLOS Biol. 14, e2000744 (2016).

82. M. Suárez-Menéndez, et al., Wild pedigrees inform mutation rates and historic abundance in baleen whales. Science 381, 990–995 (2023).

83. T. B. Wooldridge, et al., Direct measurement of the mutation rate and its evolutionary consequences in a critically endangered mollusk. [Preprint] (2024). Available at: https://www.biorxiv.org/content/10.1101/2024.09.16.613283v1 [Accessed 7 November 2024].

84. A. P. Martin, S. R. Palumbi, Body size, metabolic rate, generation time, and the molecular clock. Proc. Natl. Acad. Sci. 90, 4087–4091 (1993).

85. M. Bosse, H.-J. Megens, M. F. L. Derks, Á. M. R. de Cara, M. A. M. Groenen, Deleterious alleles in the context of domestication, inbreeding, and selection. Evol. Appl. 12, 6–17 (2019).

86. D. Orme, et al., CAPER: comparative analyses of phylogenetics and evolution in R. Methods Ecol. Evol. 3, 145–151 (2013).

87. M. Pagel, Inferring the historical patterns of biological evolution. Nature 401, 877–884 (1999).

